# Vocal repertoire of adult domestic pigs in a laboratory environment

**DOI:** 10.64898/2026.03.24.713989

**Authors:** Kathryn Y. Henley, Aimee L. Bozeman, Betty M. Pat, Candace L. Floyd

## Abstract

The use of domestic pigs in clinical training and biomedical research is expanding rapidly, increasing the need for reliable, noninvasive indicators of health and welfare. Vocal analysis offers a non-invasive promising tool, yet the acoustic repertoire of adult domestic pigs remains poorly defined. However, the vocalization repertoire of adult domestic pigs has yet to be characterized. This study characterizes the vocal repertoire of adult pigs housed in a biomedical research laboratory. Twelve mixed-breed pigs (2–3 months old; 5 males, 7 females) were recorded during routine husbandry and experimental procedures. Vocal classification was conducted using perceptual and objective clustering techniques. First, aural– visual (AV) inspection of spectrograms was used to construct a hierarchical repertoire. Second, a two-step cluster analysis based on six acoustic parameters (5% frequency, first quartile frequency, center frequency, 90% bandwidth, interquartile range bandwidth, and 90% duration) provided an objective classification. Agreement between methods was evaluated using Cramer’s V. A total of 1,136 vocalizations from 69 recordings were analyzed. AV classification revealed five major vocal classes— grunt, squeal, complex, scream, and bark—subdividing into 16 distinct call types. Standardized definitions integrating descriptive and quantitative criteria are provided. The two-step cluster analysis identified two clusters as the optimal statistical solution, with moderate agreement between methods (Cramer’s V = 0.67, p < 0.0001). Most AV-defined call types aligned with previously reported repertoires, although whines, yelps, and stable screams were unique to this study. While two-cluster solutions are commonly reported, our findings indicate that richer acoustic structure exists and that high gradation among pig calls may limit the resolution of statistical clustering. These results establish a detailed acoustic framework for adult pig vocalizations and provide essential groundwork for developing predictive models to enhance welfare assessment and support comparative research in laboratory-housed pigs.

## Introduction

The use of pigs as a model of human pathophysiology in biomedical research has increased eightfold in the past thirty years (1-3). Although multiple pig breeds are available for research, domestic farm pigs (Sus scrofa domesticus) remain substantially more cost-effective than specialized micro-or mini-pig lines. Their outbred genetic backgrounds also mirror the heterogeneity of the human population, enhancing their translational relevance (4). As the utilization of domestic pigs in laboratory-based biomedical studies continues to grow, robust and valid methods for assessing their health and welfare are urgently needed (5). Behavioral outcomes are increasingly recognized as critical endpoints in translational research, yet the very traits that make pigs valuable models such as high intelligence and cognitive complexity (3, 6) can also complicate the interpretation of behavioral responses. A clear understanding of domestic pig behavior within laboratory environments is therefore essential for accurate welfare assessment and for detecting behaviorally mediated effects of experimental conditions.

For decades, ethological and ecological studies have used vocalizations of species to identify an animal’s physical state, overall well-being, sociality, and for communication (7). Many species use vocalizations to communicate with both conspecifics and heterospecifics, and pigs are no exception (Garcia & Favaro, 2017). Pigs are socially complex animals that use vocalizations to convey biologically meaningful information to conspecifics (8, 9). Vocalizations are considered “honest signals” that accurately reflect the inner state of the animal. Changes in vocal parameters have been associated with changes physiological variables such as heart rate (10) and stress hormone levels (11, 12), and vary in response to physical pain (13, 14) and social stressors (15-17). Differences in pig vocalizations can also convey information pertaining to emotions (18) and personality types (19).

The utility of acoustic data as an outcome measure is contingent upon the characterization of the species’ vocal repertoires. The first comprehensive study of domestic pig vocalizations reported several call types in the repertoire of adult pigs and piglets (20). However, the calls were identified by perceptual analysis only within the context of emission, indicating that the classification of calls could have been biased by the situation in which they were produced. More recently, digital recording techniques and objective statistical analyses have been used to characterize the vocal repertoire of domestic pigs in an agricultural environment (21-23) and wild boars living on a nature preserve (24). Although these studies have characterized vocalizations in more naturalistic environments, there is evidence suggesting that the vocal repertoire of adult pigs in a biomedical laboratory may differ from the pigs in these studies. First, vocal parameters are affected by age (25, 26) and body weight (27), which may result in differences between adult and piglet repertoires. Second, evidence from multiple species demonstrates that vocalizations can vary with the physical characteristics of the habitat (28). For example, wild baboons produce grunts longer in duration and at higher rates in the forest compared to open areas, which may reflect an acoustic adaptation to a denser atmosphere (29). In contrast to the natural habitat of wild boars and the open, barn-like buildings of agricultural research facilities, pigs in biomedical laboratories are typically housed indoors in rooms made primarily of concrete. The reverberation and echo of the laboratory housing facilities may result in vocal adaptations in the domestic pig repertoire that are not present in pigs occupying other environments. Finally, significant differences have been reported in the duration and frequency of vocalizations during pig-to-pig interactions and human-to-pig interactions during agricultural husbandry procedures (30). However, the effect of daily human-animal interactions in a laboratory context, and if these effects change as the animals acclimate to the investigators and care staff, is currently unknown.

The objective of this study was to expand upon the existing classifications of piglet and wild boar vocalizations by characterizing the vocal repertoire of adult domestic pigs in a biomedical research laboratory. Given that pigs are a growing translational biomedical model, it is important to understand vocalizations in a laboratory environment, as well as how they change in certain contexts. We recorded male and female farm pigs in a variety of contexts, including interactions with humans and conspecifics, both before and after an acclimation period. We then used perceptual and objective clustering techniques to create a hierarchical classification of call types. Further, there is a need for standardized definitions of call types to create a common language among researchers and allow accurate comparisons across laboratories. To this end, we coupled descriptive labels of each call type (e.g., grunts, squeals) with aural-visual descriptions and defining acoustic characteristics.

## 2. Materials and Methods

### 2.1. Animals

Twelve male (n=5) and female (n=7) mixed-breed domestic pigs were obtained from Snyder farms (males) and Auburn University (females) at two months of age and a weight of 11-13 kg. All animals were housed singly in 1.4 m x 2.3 m modular galvanized stainless steel pens with elevated floors coated in plastisol. Visual and physical contact with pigs in neighboring pens was maintained and plastic toys and metal chains were provided for additional enrichment. The pens were located in a single room in an indoor facility maintained at 72±2°F, 30-70% relative humidity, and on a 12-hour light/dark cycle (6:00 AM to 6:00 PM). The facility contained an HVAC system that provided at least 10 air changes per hour of un-recirculated air. The pigs had access to cold tap water *ad libitum* and were fed 500-600 g of standard laboratory pig feed (SoBran Bioscience, Inc., Fairfax, VA, USA) twice daily at 6:00 AM and 1:00 PM. All procedures were approved by the Institutional Animal Care and Use Committee of the University of Alabama at Birmingham and conducted in accordance with the National Institutes of Health guidelines.

### 2.2. Experimental design and habituation

Figure 1 provides an overview of the experimental design. Pigs were recorded upon their arrival prior to familiarization with the environment and subsequently completed our laboratory acclimation protocol. Animals were allowed one week to habituate to their pens, pen mates, and facility schedule. After this acclimation period, investigators approached the animals in their home pens and encouraged interaction using treats and other positive reinforcement. Once an animal remained calm when approached and touched on the head and snout, a nylon slip leash was placed loosely around the torso of the animal. The tension of the leash was slowly increased with gentle tugs by the investigators. Animals were considered acclimated when they could be led calmly around the pen with the leash, which typically occurred within 2-3 days after leash introduction. Post-acclimation recording, which consisted of the same context as pre-acclimation recording, was then performed.

**Figure 1.**
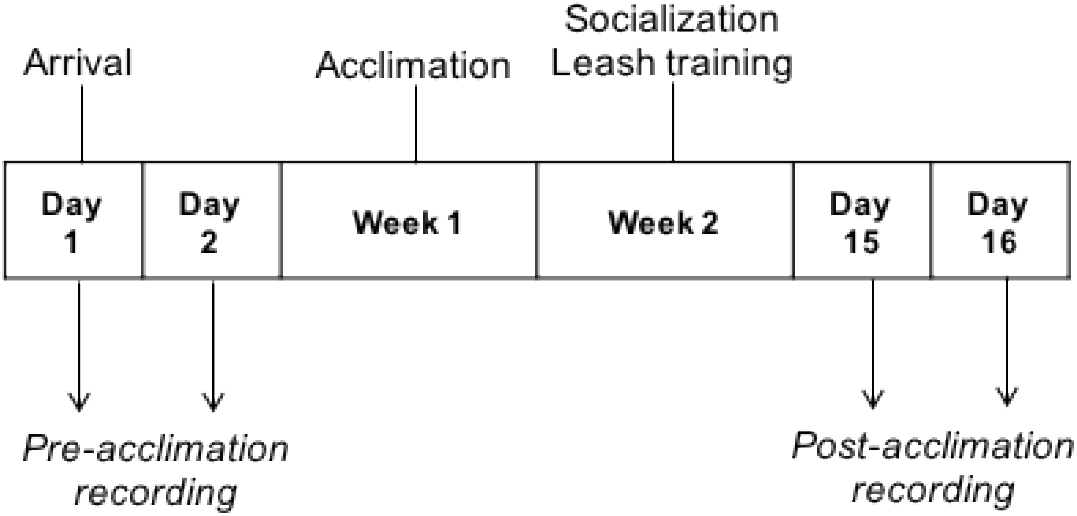
Overview of experimental design. Male and female farm pigs were recorded during typical biomedical laboratory procedures on the day of (Day 1) and day after (Day 2) their arrival. These recordings served as the pre-acclimation vocalizations. Over the next two weeks, all animals underwent our standard laboratory acclimation protocol. Animals were habituated to their cages and cage mates for seven days, followed by increasing levels of investigator contact and leash training. Post-acclimation recordings were then conducted during the same laboratory procedures.

### 2.3 Vocalization recordings

Pigs were actively recorded using a beyerdynamic MCE 86 S II CAM electret condenser shotgun hypercardioid microphone (beyerdynamic Inc., Farmingdale, NY, USA) and a GoPro® Hero 2 camera (GoPro, Inc., San Mateo, CA, USA) at a sampling rate of 48 kHz and a 16-bit sample size. One investigator maintained the position of the microphone approximately 60 cm in front of the animal with a tripod boom microphone stand while a second investigator applied the stimulus or procedure (Figure 2). As displayed in Table 1, recording contexts were split into two categories: laboratory procedures and social interactions. Laboratory procedures consisted of events initiated by the investigators that occurred during routine husbandry practices or the acclimation protocol. During investigator approach, one person approached the home pen, entered, and walked slowly toward the animal. Once in the home pen the investigator gently stroked the animal on the back and head, which served as the investigator touch stimulus. During board constraint, the animal was contained in one area of the pen with a large herding board. The intramuscular injection consisted of a 21G needle inserted in the area between the shoulder blades to simulate medication administration. During food presentation, animals were recorded while an investigator distributed food during the 1:00 PM feeding. During leashed walks, animals were recorded as they were led out of the room and into the hallway. In contrast, social situations were comprised of interactions between two or more animals. These included de-crating, during which animals entered the room in their transportation crates and were released into their home pens. Animals were also recorded while being separated and reintroduced to their pen mates when one was taken from the room. Finally, social interactions were recorded during establishment of the social hierarchy.

**Table 1.**
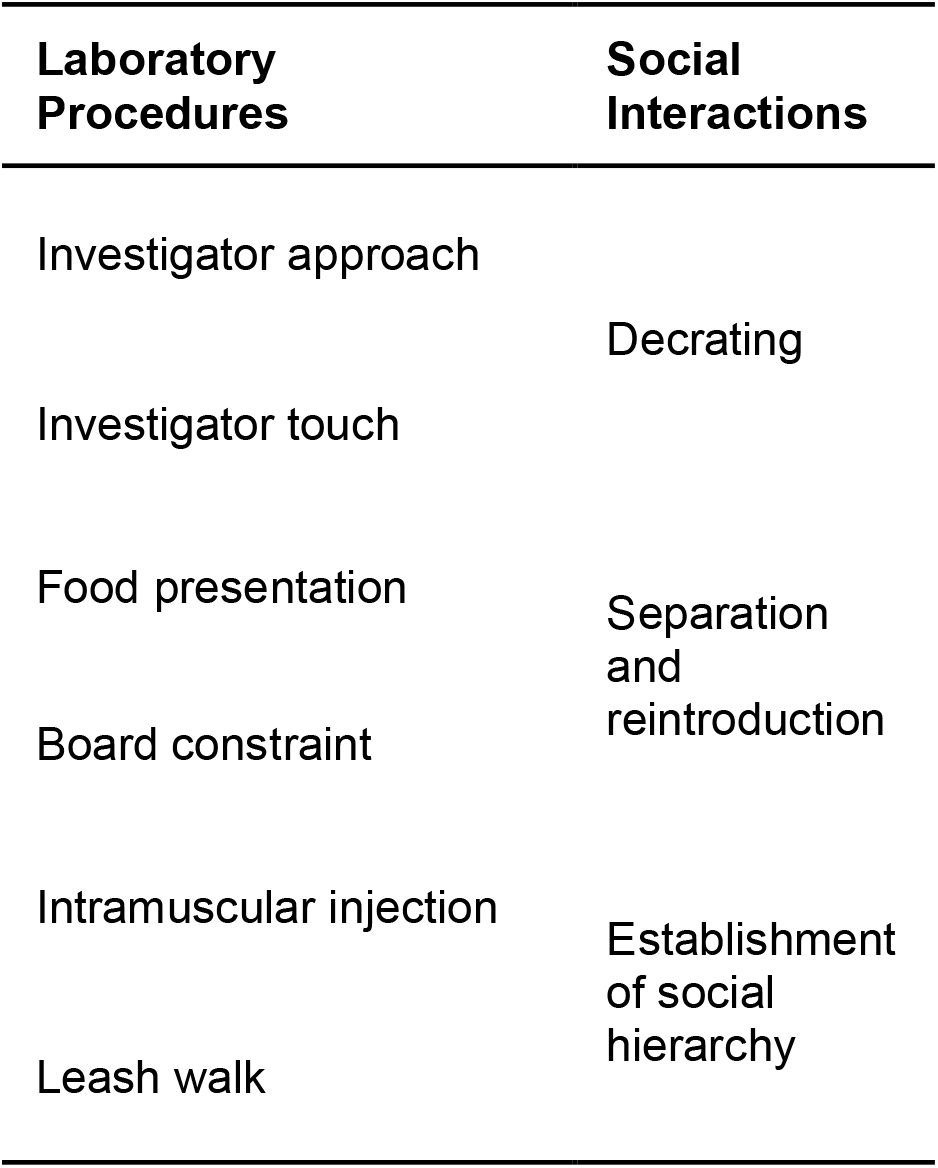
Recording situations. Animals were recorded during procedures typical to a biomedical research facility, including interactions with both humans (laboratory procedures) and conspecifics (social interactions). All recordings took place in the home pen except for the leash walk, which was recorded as the animal was led out of the room and into the hallway of the facility.

**Figure 2.**
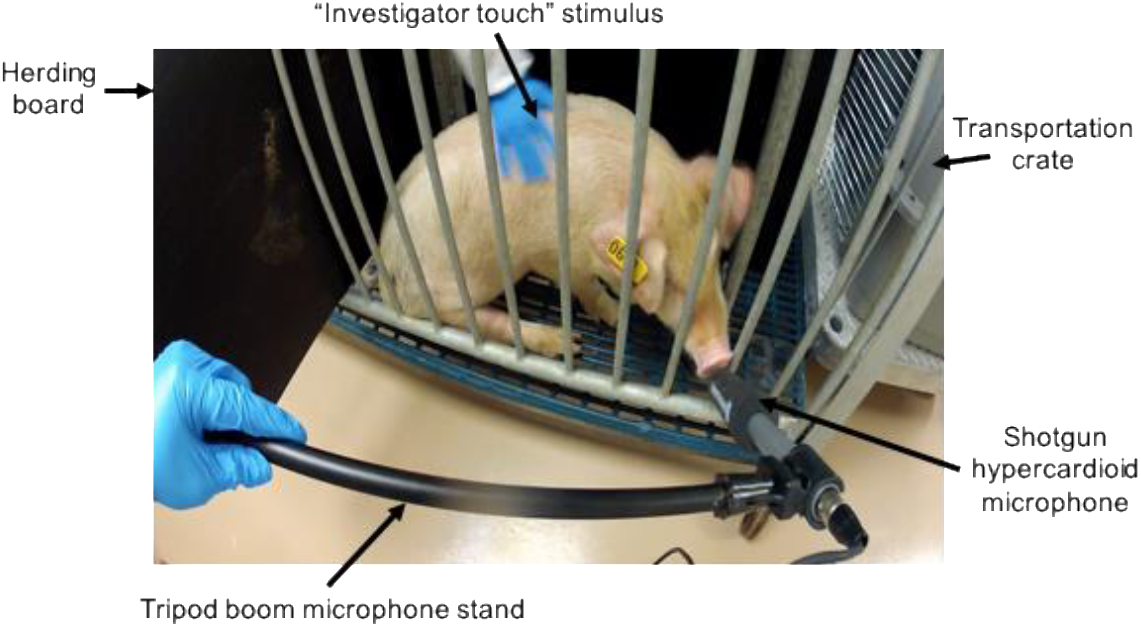
Recording set-up in the home pen. One investigator maintained the microphone (beyerdynamic MCE 86 S II CAM electret condenser shotgun hypercardioid microphone, beyerdynamic Inc., Farmingdale, NY, USA) in front of the animal using a tripod boom microphone stand while another investigator applied the stimulus. A GoPro® Hero 2 camera (GoPro, Inc., San Mateo, CA, USA) was used for video recording.

### 2.4 Vocal classification and statistical analysis

Spectrograms were generated from.WAV audio files using Raven Pro 1.5 (RRID: SCR_028144) with a 2048-point discrete Fourier transform (DFT), Hann window with 23.4 Hz resolution, and 50% overlap between consecutive analysis windows. Vocalizations with start and end points that could be aurally and visually distinguished were identified by an investigator (JP) naïve to the situation and sex of the animal. Vocalizations were excluded from analysis if they overlapped with other sounds (e.g., anthropogenic or facility noises) or had excessive amounts of echo such that the start or end points could not be identified. Duration and frequency of vocalizations for selected calls were calculated by Raven Pro 1.5. To reduce the potential effects of manual call selection on measured values, only robust measures were calculated. These measures represent center or percentage calculations that are minimally affected by manual selection compared to maximum or peak measures. As detailed in Table 2, these included measures of frequency: 5% frequency (f5), 95% frequency (f95), first quartile frequency (Q1f), third quartile frequency (Q3f), and center frequency (Cf); frequency modulation: 90% bandwidth (BW90) and interquartile range bandwidth (IQRBW); and duration: 90% duration (DUR90).

**Table 2.** Measures calculated for each vocalization. Robust measures were chosen to reduce the potential influence of manual selection of the vocalizations. All measures were calculated by Raven Pro 1.5 sound analysis software (Bioacoustics Research Program, Cornell Laboratory of Ornithology, Cornell University, Ithaca, NY, USA). Center and percentage calculations of both duration and frequency were utilized. Abbr: abbreviation; Q1: first quartile; Q3: third quartile; IQR: interquartile range; Hz: hertz; s: seconds.

| Measure | Units | Abbr. | Description |
| --- | --- | --- | --- |
| 5% Frequency | Hz | f5 | Frequency that divides the energy into 5% and 95% |
| 95% Frequency | Hz | f95 | Frequency that divides the energy into 95% and 5% |
| 90% Bandwidth | Hz | BW90 | Difference between 5% and 95% frequencies |
| Q1 Frequency | Hz | Q1f | Frequency that divides the energy into 25% and 75% |
| Q3 Frequency | Hz | Q3f | Frequency that divides the energy into 75% and 25% |
| IQR Bandwidth | Hz | IQRBW | Difference between Q1 and Q3 |
| Center Frequency | Hz | Cf | Frequency that divides the energy in half |
| 90% Duration | s | DUR90 | Difference between 5% and 95% times |

#### 2.4.1 Aural-visual classification

To classify the vocalizations, we utilized two approaches: an aural-visual (AV) classification based on sound and spectrographic characteristics, and a two-step cluster analysis. The AV analysis followed previous methods used to classify humpback whale calls (31). Briefly, an investigator (KH) naïve to the situation and sex of the animal(s) recorded examined each vocalization from the sound files presented in a randomized order. Based off the descriptions of pig vocalizations reported in previous studies (11, 20, 21, 24), the vocalizations were first broadly grouped into vocal classes. The vocalizations within each vocal class were then randomized and examined for differentiating characteristics to create vocal subclasses. Finally, all vocalizations within the vocal subclasses were randomized and examined for further grouping characteristics to create call types. To reduce the effect of individual variation, only vocalizations that were emitted by more than one animal were included. Although previous work was used to guide the perceptual analysis, the number of classes, subclasses, and call types was not pre-determined.

Sound measure values for each vocalization were imported into IBM SPSS Statistics (IBM, Armonk, NY) for all subsequent analyses. First, box plots were created to examine outliers in each call type. Vocalizations with any sound measure value greater than 1.5x the interquartile range away from the 25^th^ or 75^th^ percentile were reexamined on the spectrogram. The vocalization was either reclassified, retained in the same classification, or, if the signal-to-noise ratio was less than 10 decibels (dB), excluded from analysis (31). Finally, the average value of each sound measure was calculated to provide objective descriptors of vocal classes and call types to accompany the AV descriptions.

### 2.4.2 Cluster analysis

A two-step cluster analysis was performed to objectively classify the vocalization data set. The two-step clustering technique first creates pre-clusters using a modified cluster feature tree and then these pre-clusters are clustered using agglomerative hierarchical methods (32). First, sound measure values were transformed to z-scores to obtain unit-free variables, which were used in all subsequent analyses. Sound measures were then examined for multicollinearity using a Pearson’s correlation matrix. If the pair-wise correlation coefficient between two measures exceeded 0.90, one was excluded from the analysis (21). Out of the two measures, the one with the highest correlation with other measures was chosen for exclusion. These remaining measures used for the cluster analysis were: f5, BW90, Q1f, IQRBW, Cf, and DUR90. A two-step cluster analysis was performed using Euclidian distances (21, 24), a clustering range of 1-20 clusters, and Schwarz’s Bayesian information criterion (BIC; (33). The optimal number of clusters was selected by the two-step algorithm by considering the BIC value and ratio of distance measures.

#### 2.4.4 Agreement between AV classification and unsupervised clustering

The relationship between the AV classification results and optimal two-step cluster solution was assessed using Cramer’s V. The cluster membership variable from the two classification techniques were used to create a Chi-square contingency table, from which Cramer’s V is computed using the Chi-square test statistic. Significance criterion was *p* ≤ 0.05.

#### 2.4.6 Effect of non-independence

Independent responses are preferred when inferring results to a population. This assumption is violated in the present study, as each pig contributed multiple vocalizations to the data set. A mixed linear model was conducted to determine the effect of individual pigs on the results from the AV and two-step cluster analysis. Cluster membership served as the independent variable and individual pigs were entered as the random effect. Cf was chosen as the dependent variable due to its high predictor importance in the two-step cluster results.

## 3. Results

### 3.1 Aural-visual classification structure

A total of 1174 vocalizations were manually identified from 71 recordings of 12 animals. After outliers were excluded, a total of 1136 high-quality vocalizations from 69 recordings of 12 animals remained. All vocalizations were grouped based on their aural and visual characteristics into a total of 5 vocal classes, 11 subclasses, and 16 call types. As displayed in Figure 3, vocal classes included grunts, squeals, screams, complex vocalizations, and barks. All vocal classes except for barks were divided into subclasses. Grunts were separated into classic and short subclasses. Within the classic grunt subclass, call types croak, growl, and modulated were identified. Squeal subclasses included classic, yelp, whine, and squeak. Complex vocalizations were divided into complex screams and complex squeals, which were separated into complex yelp and complex whine call types. Scream subclasses included modulated and stable.

**Figure 3.**
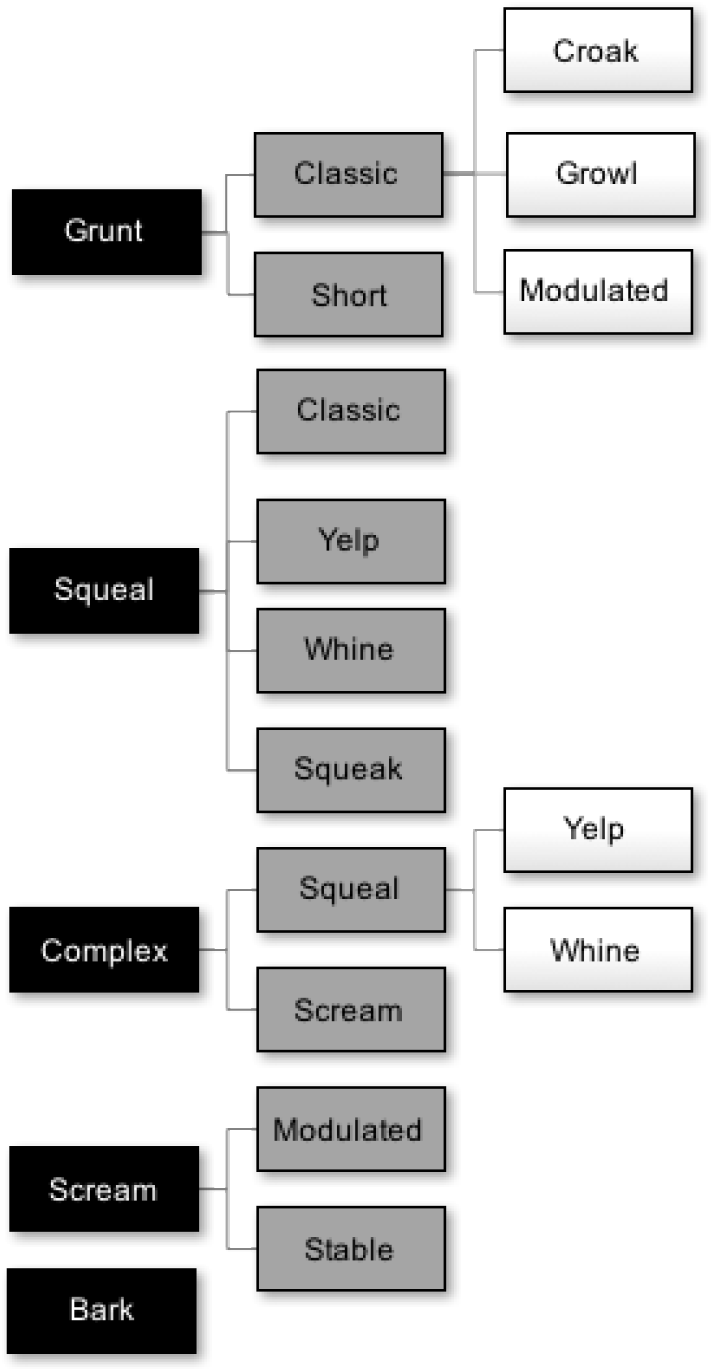
Hierarchical vocalization repertoire structure identified by aural-visual (AV) analysis. A total of 1136 vocalizations were grouped and labeled based on AV characteristics, independent of the situation or context in which the vocalization was emitted. Five vocal classes were initially identified. All vocal classes (except for barks) were further separated into vocal subclasses. Classic grunts and complex squeals were further divided into call types. A total of 16 distinct call types were identified.

Figure 4 shows the distribution of each vocal class (Figure 4a) and call type (Figures 4b-e). Grunts were the most common, comprising 81.95% of all vocalizations. Squeals were the next most common, followed by complex vocalizations, screams, and barks. The classic grunt was the most common call type, comprising 66.46% of all vocalizations. Classic squeals and complex squeals were the most common call type within their vocal classes, comprising 43.27% of squeals and 44.83% of complex vocalizations, respectively. Modulated screams and stable screams were distributed fairly equally within their vocal class.

**Figure 4.**
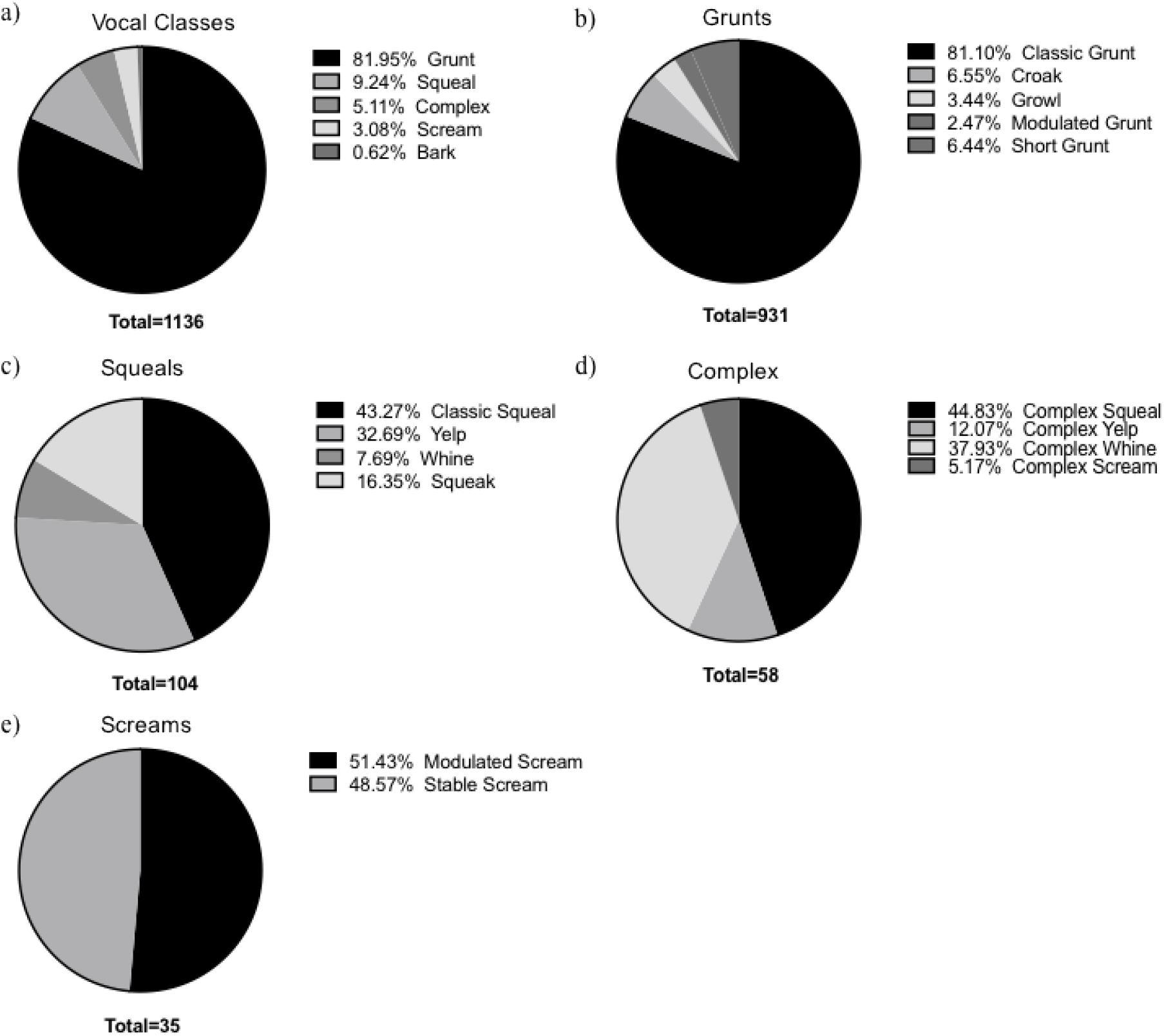
Distribution of vocalizations by class and call type. The percentage of vocalizations comprising each vocal class (a) and with each call type (b-e) are presented above. All 1136 vocalizations were classified into 1 of 16 call types identified in AV analysis.

#### 3.1.2 Aural-visual descriptions and defining measures

Spectrograms of representative vocalizations from each call type are provided in Figure 5. Grunts exhibited a pulsatile structure and a low-frequency, nasally produced sound. Squeals had a high degree of noise, as indicated by bursts of energy on the spectrogram at high frequencies and of short durations.

**Figure 5.**
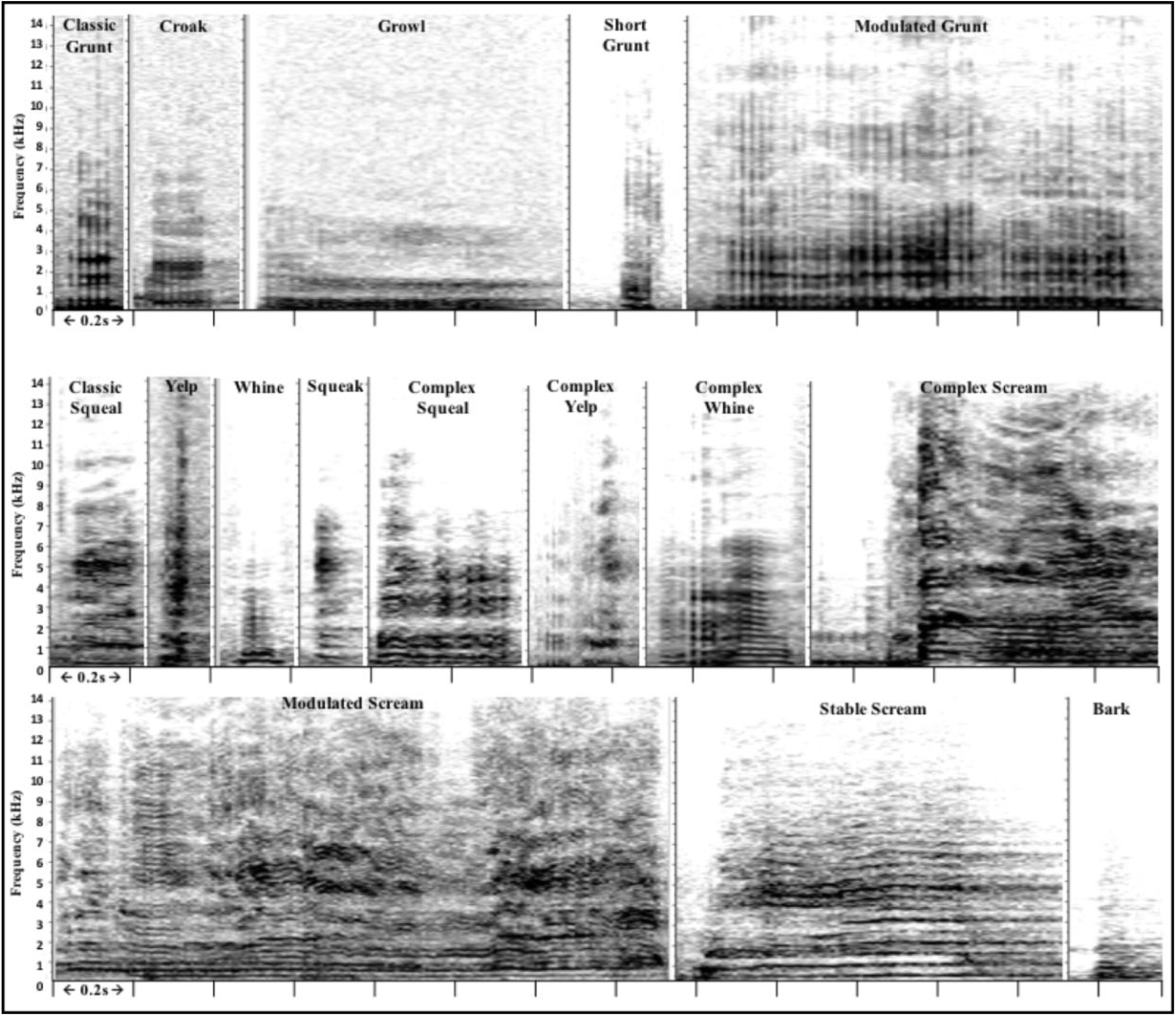
Representative spectrograms of each call type. Aural-visual analysis identified 16 distinct call types in 1136 vocalizations from 12 domestic pigs. Differences can be observed in duration, frequency levels, and frequency modulation in addition to structural characteristics. Time is on the x-axis with 0.2 seconds (s) between each tick mark. kHz: kilohertz.

Whines were identified by the presence of short-duration harmonics. Complex vocalizations consisted of vocalizations from separate vocal classes emitted in the same call, either in succession or a blended format. Typically, complex vocalizations began as a grunt and transitioned into a squeal. Screams were long in duration and high in frequency. Modulated screams consisted of harmonics with a tonal quality and high degree of frequency modulation, interspersed with periods of noise. In contrast, stable screams showed harmonics with little frequency modulation or noise. Barks were short bursts of energy at low frequencies.

The average sound measures for each call type are displayed in Figures 6a-b. Of all the call types, screams had the highest frequency, frequency modulation, and duration. In general, grunts and barks had the lowest frequency and shortest duration. Squeals and complex vocalizations had moderate levels of frequency, frequency modulation, and duration. The defining measures and AV descriptions for each call type are displayed in Table 3. A measure was considered “defining” if the average value was considerably higher or lower than that of other call types based on visual inspection of the graphs in Figures 6a-b.

**Table 3.** Aural and visual (AV) descriptions and defining measures. Each call type is described by its aural characteristics and visual presentation on the spectrogram. Sound measures that differentiate the call types are listed. All measures are in hertz except for DUR90 (s). The descriptive terms (e.g., long, high, etc.) that accompany the measures are relative to the other call types. Sound measures that define the vocal classes are also listed. The descriptive terms are relative to other vocal classes. f: frequency; dur: duration; DUR90: 90% duration; f5: 5% frequency; f95: 95% frequency; BW90: 90% bandwidth; Q1f: quarter 1 frequency; Q3f: quarter 3 frequency; IQRBW: interquartile range bandwidth; Hz: hertz; s: seconds.

| Call Type | n | AV Description | Call Type<br>Defining Measures | Vocal Class<br>Defining Measures |
| --- | --- | --- | --- | --- |
| **Classic Grunt** | 754 | Pulsed, vibrating, low nasal snort. | Short DUR90 = 0.12 | **Grunts**<br>Low f5 = 141.42<br>Low Q1f = 270<br>Low DUR90 = 0.12 |
| **Croak** | 61 | Pulsed, vibrating, smacking sound, higher f than classic grunt. | Short DUR90 = 0.11<br>High f5 = 300.00 |  |
| **Growl** | 31 | Pulsed, vibrating, long dur, lower f than classic grunt. | Long DUR90 = 0.40<br>Low BW90 = 703.83 |  |
| **Modulated** | 23 | Pulsed, harmonic, vibrating, slight f modulation. | Long DUR90 = 0.39 |  |
| **Short Grunt** | 60 | Wide/short energy burst, noisy, may be pulsed, no vibration, high or low f. | Short DUR90 = 0.07 |  |
| **Classic Squeal** | 45 | Wide energy burst, noisy, high f, no modulation. | Long DUR90 = 0.28 | **Squeals**<br>Short DUR 90 = 0.18<br>High BW90 = 4471.88 |
| **Yelp** | 34 | Narrow/tall energy burst, high f, short dur, no modulation. | Best distinguished by AV analysis |  |
| **Whine** | 8 | Harmonic, low f, short dur, emitted upon exhale. | Low Q1f = 251.94<br>Low BW90 = 1163.09 |  |
| **Squeak** | 17 | Narrow/tall energy burst, high f. Some harmonics. Higher f than whine and yelp. | High Q1f = 1255.97<br>High IQRBW = 2732.54 |  |
| **Complex Squeal** | 26 | Pulsed with noise. Grunt followed by squeal or blend of the two. | High Q1f = 659.85 | **Complex**<br>Moderate<br>DUR90 = 0.45<br>Moderate<br>BW90 = 3939.93 |
| **Complex Yelp** | 7 | Pulsed followed by narrow/tall energy burst. Grunt followed by yelp. | Short DUR90 = 0.20 |  |
| **Complex Whine** | 22 | Pulsed followed by harmonics. Grunt followed by whine. | Low Q1f = 381.39<br>Low BW90 = 3435.74 |  |
| **Complex Scream** | 3 | Pulsed followed by long/tall energy burst. Grunt followed by scream. | Long DUR90 = 0.83<br>High BW90 = 5898.47 |  |
| **Modulated Scream** | 18 | Long/tall energy burst, noisy, harmonic, long dur, high f, high f modulation. | Long DUR90 = 0.98<br>High f5 = 546.89<br>High IQRBW = 3377.60 | **Screams**<br>High f5 = 485.49<br>High BW90 = 5518.51<br>High Q1f = 1131.03<br>High IQRBW = 2872.10<br>Long DUR 90 = 0.87 |
| **Stable Scream** | 17 | Long/tall energy burst, noisy, harmonic, long dur, high f, low f modulation. | Short DUR90 = 0.75<br>Low f5 = 420.48<br>Low IQRBW = 2336.86 |  |
| **Bark** | 7 | Short energy burst, low f. Burst of sound. | Short DUR90 = 0.17<br>Low Cf = 592.64<br>Low BW90 = 1094.9 | **Barks**<br>Low Q3f = 723.23<br>Low IQRBW = 311.40 |

**Figure 6a.**
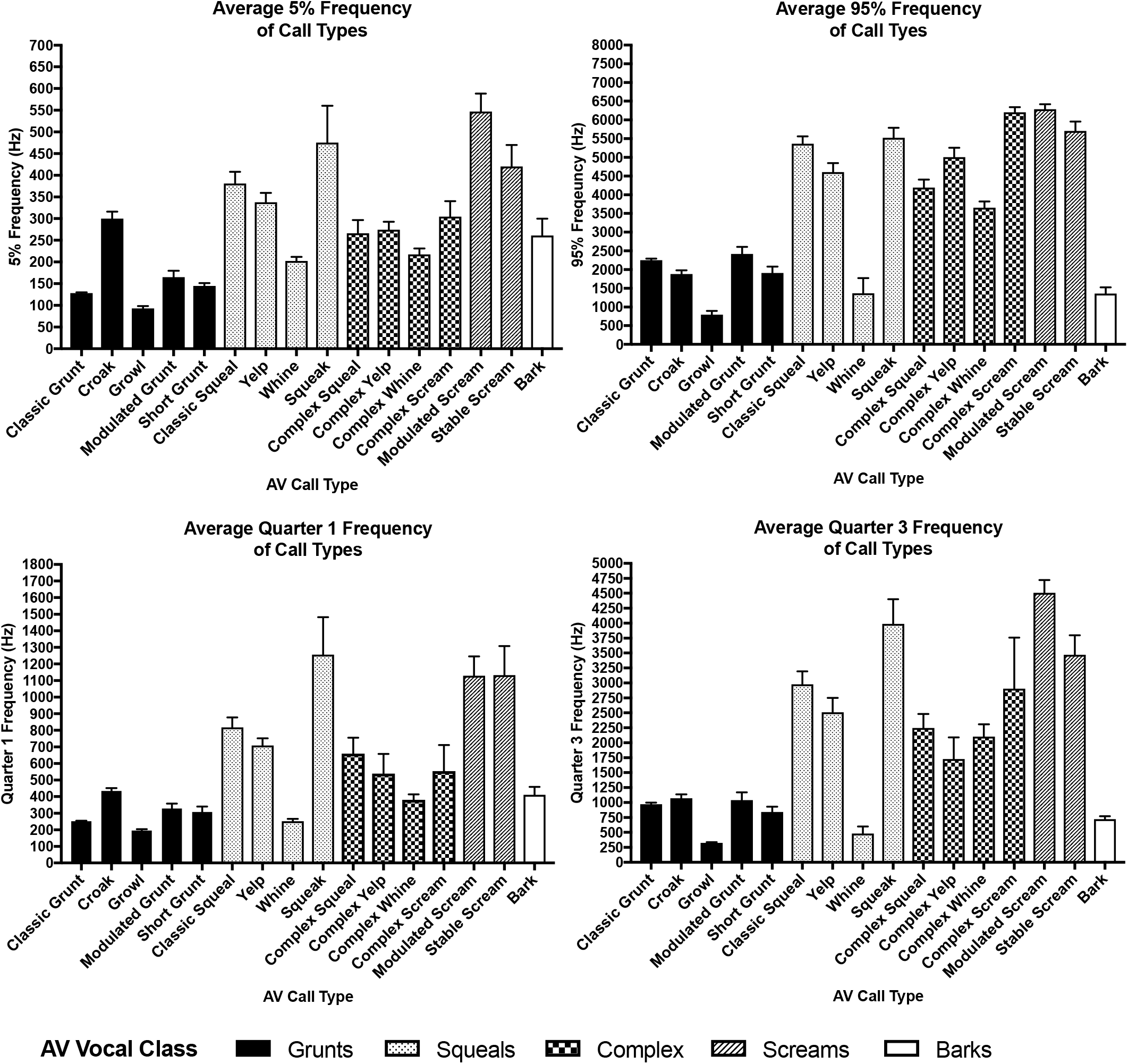
Average frequency sound measures for call types. In general, screams, squeals, and complex vocalizations had the highest frequency measures, and grunts and barks had the lowest.

**Figure 6b.**
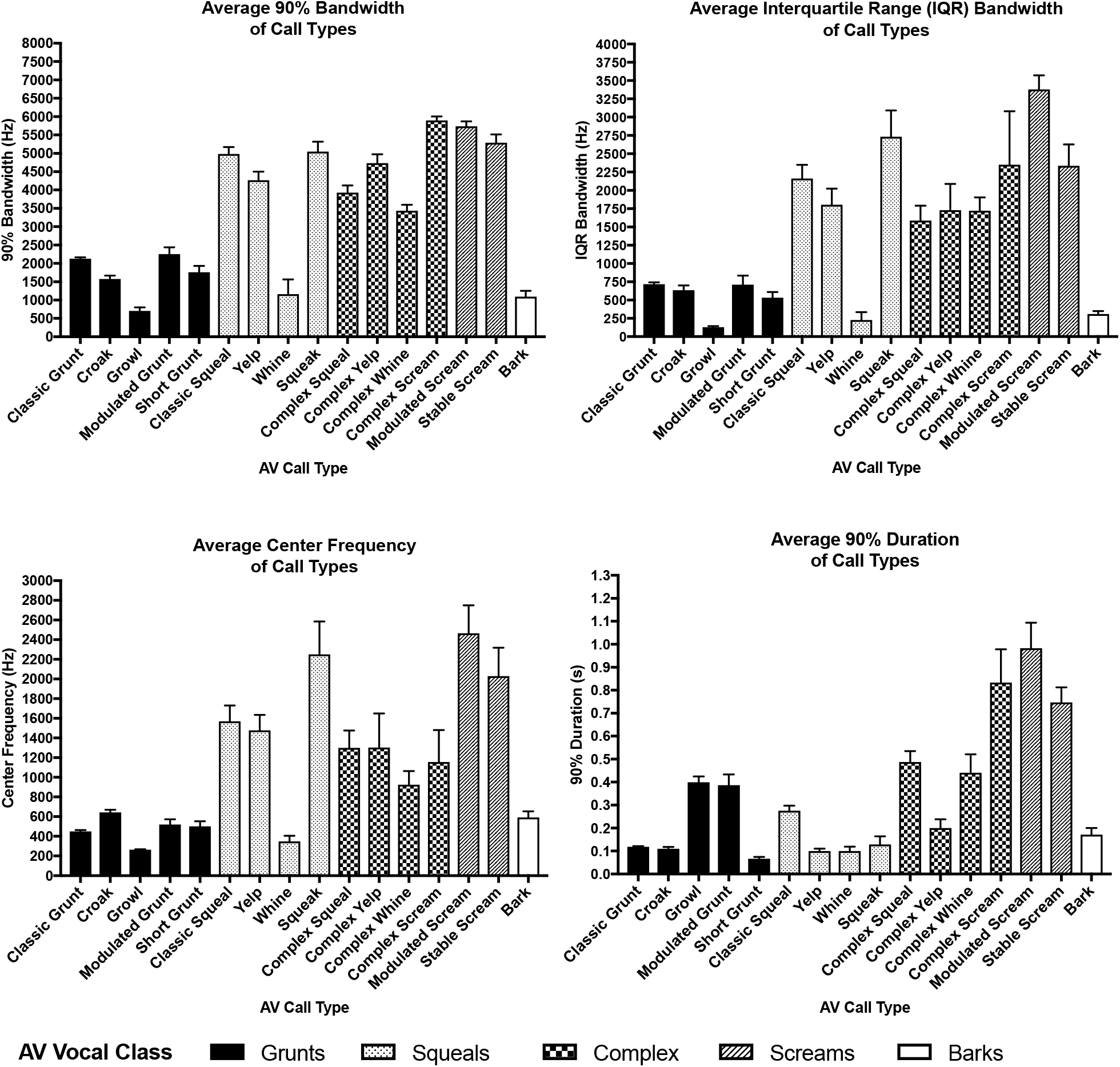
Average frequency modulation, center frequency, and duration sound measures for call types. Typically, screams and complex vocalizations had the highest frequency modulation as measured by 90% bandwidth and interquartile range bandwidth. Grunts and barks had the lowest. Screams and complex vocals had the longest duration whereas grunts, squeals, and barks had the shortest.

### 3.2 Cluster analysis

Two-step cluster analysis was performed to objectively classify the data. The clustering range was set at 1-20 to accommodate the number of groups identified in the AV analysis (i.e., 5 vocal classes, 11 subclasses, and 16 call types). The BIC values for the clustering range are displayed in Table 4. The analysis revealed two clusters as the optimum solution with a BIC value of 4325.82, ratio of distance measures of 1.15, and good cluster quality. There are also decreases in BIC value at 5 and 19 clusters. The most important predictor for cluster membership was Cf (1.0), followed by Q1f (0.96), f5 (0.77), DUR90 (0.59), IQRBW (0.39) and BW90 (0.33). Cluster 1 comprised 95.90% of cases and was associated with lower values in all sound measure variables compared to Cluster 2. The clusters are displayed by Cf and Q1f in Figure 7.

**Table 4.** Two-step cluster results. Schwarz’s Bayesian information criterion (BIC) and ratio of distance measures from a two-step cluster analysis ranging from 1-20 clusters. The analysis selected two clusters as the optimal solution based on its low Schwarz’s Bayesian information criterion (4325.82) and high ratio of distance measures (1.15) compared to other solutions. There are also marked decreases in BIC at five and 19 clusters. ^a^ The changes are from the previous number of clusters in the table. ^b^ The ratios of changes are relative to the change for the two cluster solution. ^c^ The ratios of distance measures are based on the current number of clusters against the previous number of clusters.

| # of Clusters | Schwarz's BIC | BIC Change<sup>a</sup> | Ratio of BIC Changes<sup>b</sup> | Ratio of Distance Measures<sup>c</sup> |
| --- | --- | --- | --- | --- |
| 1 | 4809.735 |  |  |  |
| 2 | 4325.818 | -483.916 | 1 | 1.153 |
| 3 | 4373.364 | 47.546 | -0.098 | 1.629 |
| 4 | 4443.912 | 70.548 | -0.146 | 1.716 |
| 5 | 3037.092 | -1406.821 | 2.907 | 1.001 |
| 6 | 3101.292 | 64.201 | -0.133 | 1.305 |
| 7 | 3177.053 | 75.761 | -0.157 | 1.133 |
| 8 | 3252.307 | 75.254 | -0.156 | 1.076 |
| 9 | 3329.279 | 76.972 | -0.159 | 0.922 |
| 10 | 3331.456 | 2.177 | -0.004 | 1.023 |
| 11 | 3406.401 | 74.944 | -0.155 | 1.244 |
| 12 | 3427.435 | 21.034 | -0.043 | 1.031 |
| 13 | 3469.195 | 41.76 | -0.086 | 1.18 |
| 14 | 3547.935 | 78.739 | -0.163 | 0.951 |
| 15 | 3628.439 | 80.504 | -0.166 | 1.064 |
| 16 | 3706.084 | 77.645 | -0.16 | 1.004 |
| 17 | 3773.826 | 67.742 | -0.14 | 1.06 |
| 18 | 3800.389 | 26.563 | -0.055 | 1.034 |
| 19 | 3627.983 | -172.406 | 0.356 | 1.114 |
| 20 | 3706.711 | 78.728 | -0.163 | 1.017 |

**Figure 7.**
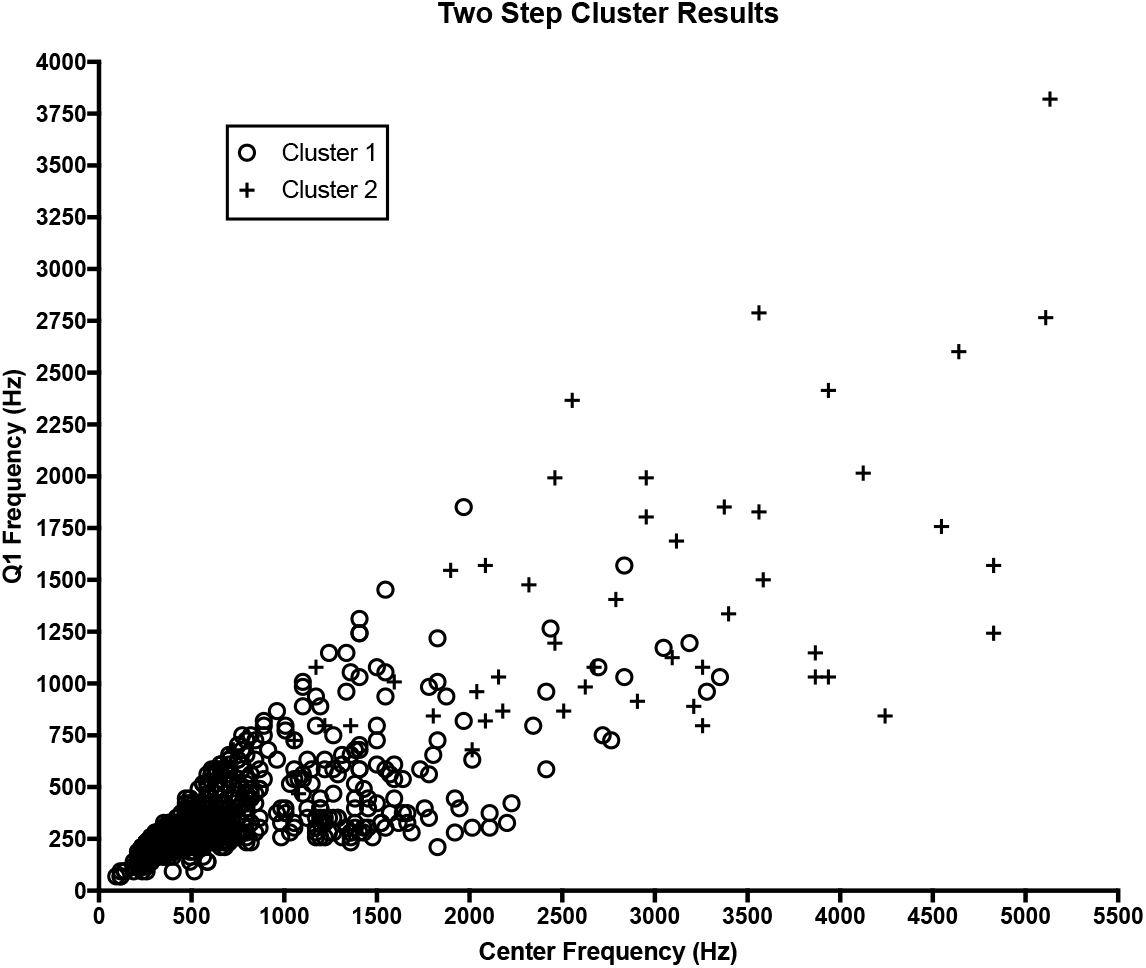
Two-step cluster analysis results. Each vocalization is displayed above according to center frequency and Q1 frequency. The majority of vocalizations (95.9%) were assigned to Cluster 1, which was defined by lower frequencies compared to Cluster 2. There is considerable overlap between clusters at moderate frequency levels. Center frequency and Q1 frequency were chosen based on their high predictor importance to the two-cluster solution. Q1: quarter one.

### 3.3 Agreement between AV classification and unsupervised clustering

Cramer’s V was used to assess the agreement between the groups determined by AV analysis and two-step clustering. Cramer’s V for AV call type and the two-step cluster solution was 0.67 (p<0.0001).

### 3.5 Effect of non-independence

A mixed effects linear model was used to assess the influence of individual pigs on Cf in the cluster results. This explained 3.67% (p=0.08), 2.31% (p=0.12), and 3.24% (p=0.11) of the variance in Cf in AV vocal class, subclass, and call types, respectively. In the two-step cluster results, individual pig explained 3.04% of the variance in Cf (p=0.10).

## 4. Discussion

### 4.1 Classification of vocalizations

Given that the use of swine as a translational research model has been growing rapidly, the objective of the current study was to create a hierarchical vocalization repertoire from recordings of adult domestic pigs in situations typical to a biomedical research facility. Five distinct vocal classes were identified with AV analysis, which was then delineated into 16 total call types. Importantly, our AV analysis was performed by an investigator naïve to the situations in which the animal was recorded to decrease bias. Using a two-step cluster analysis, we found that two clusters were the optimal solution for the data set, with five and nineteen clusters as additional strong solutions. Finally, we found that individual differences in vocalizations did not significantly affect cluster formation in the AV or cluster analyses.

Grunts, squeals, grunt-squeal combinations, and screams have been frequently reported in the pig vocalization literature. The frequency, duration, and structure of these calls, which were identified in adult domestic pigs (11, 20), wild boars (24), and domestic piglets (21), correspond to our AV and objective definitions of the vocal classes. To a lesser extent, barks have also been reported (18, 20, 24) and correspond to our definition of the vocalization: abrupt onset, short duration, and low frequency.

Multiple subtypes of grunts as identified in our study have also been previously reported. The “long grunt” and “staccato grunt” formerly identified in adult domestic pigs are similar in duration and spectrographic structure to our growl or modulated grunt and short grunt, respectively (20). The croaks identified in our study correspond to the “low frequency tonal” vocalizations described by Tallet, Linhart (21), the “chirrups” identified by Kiley (20) and the “quacks” reported by Appleby, Weary (34). Interestingly, these studies included piglets only, and our study is the first to report croaks in adult pigs.

We also identified several types of squeals. However, only squeaks have been previously reported and were categorized with squeals as “high-frequency stable” vocalizations (21). We separated squeaks from classic squeals by identifying shorter durations and higher Q1f and IQRBW. Yelps have not been reported previously, most likely because they were difficult to distinguish from barks and were categorized as such in other work. However, we were able to distinguish yelps from barks because of the higher frequency characteristic of yelps. Similarly, whines have not been previously reported in pigs. While they were rare in the present study, they were easily distinguished from other call types by the presence of short-duration harmonics. Given that whines were much lower in frequency than the rest of squeal call types, they may be best categorized as a unique vocal class. Interestingly, Garcia, Gingras (24) reported “trumpets” as a vocal class in wild boars, which have not been reported in domestic pigs. Like whines, trumpets were characterized by harmonics, high frequency, and low intensity. While it is possible that trumpets are vocalizations unique to wild boars or to that specific herd, they may also be a variant of the whines we identified in our adult domestic pigs.

This is the first study to report modulated and stable screams in the same data set. Tallet, Linhart (21) reported both stable and modulated high-frequency vocalizations in piglets; however, they referred to the stable vocalizations as squeals and differentiated screams by their high degree of frequency modulation. High frequency modulation has also been used to describe screams in response to stressors in adult pigs (11). We chose to classify the stable high-frequency vocalizations in our study as screams due to their long duration compared to squeals (average DUR90 of 0.75 s compared to 0.18 s). However, the comparison of duration (and all other sound measures) between call types in the present study is strictly descriptive. Future work should focus on statistically determining cut-off points to differentiate call types.

To date, no studies have identified complex vocalizations other than grunt-squeal combinations. This is not surprising, as yelps, whines, and squeaks have not been reported or were reported as other call types (i.e., squeals or trumpets). Interestingly, although screams are ubiquitous throughout the pig vocalization literature, complex screams have not been previously reported. The complex screams in our study were rare and consisted of grunts followed by screams. Therefore, it is possible that the grunt and scream were identified as two separate vocalizations in previous work. In the present study, we considered vocalizations to be separate if there was any time delay between energy displayed on the spectrogram, which reduced the possibility of echo influencing vocal measures. In past studies, the “repeated grunt,” which includes 0.2-1.5s between individual calls, has been reported as a type of combined vocalization (20). Further work is needed to identify the differentiating characteristics of single and combined vocalizations, as well as determining their importance in pig communication.

### 4.2 Cluster analyses of vocalizations

The two-step cluster analysis revealed two clusters as the optimal solution for the vocalization data set, similar to what has been found in cluster analyses of domestic piglet (21) and wild boar vocalizations (24). Moreover, many studies that use vocalizations as a behavioral outcome measure split the calls into two groups: high frequency and low frequency (19, 35-42). Interestingly, classification of domestic piglet vocalizations showed that in addition to the two-cluster solution, five clusters was also a strong solution, followed by 10, 12, and 15 clusters (21). In the present study, we showed a peak in BIC value at 5 and 19 clusters. Taken together, these results show that while most objective clustering techniques will reveal two clusters as ideal, there is evidence that smaller clusters may naturally exist in the pig vocal repertoire. Further support for this point is described in the ‘social complexity hypothesis,’ which states that socially complex species, such as pigs, require vocal repertoires comprised of many structurally distinct components (43). There is likely more complexity in the pig repertoire than two classes constituting high and low frequency vocalizations, but certain characteristics inherent to the vocalizations may limit the ability of clustering methods to reliably classify more than two groups. For example, previous studies reported gradation in pig vocalizations, suggesting that some pig vocalizations do not fit into discrete categories but may exhibit characteristics of multiple classes or call types (20, 21, 24). Therefore, a clustering technique that allows for membership in more than one cluster, such as fuzzy c-means (44), may be more appropriate to classify pig vocals and could be utilized in future work.

The agreement between our AV and cluster analyses was high, which strengthens our perceptual classification results. Further, we showed that the effect of variability within and between pigs did not significantly influence the AV or two-step cluster classifications. However, it should be noted that only Cf was used in the mixed model and that the influence of individual variation may be apparent in other measures. Additional analysis of the repertoire presented here is needed to overcome the limitations inherent in perceptual analysis. Future work should use discriminate function analysis or regression modeling to assess the ability of the data set to predict vocalization type, as well as the context in which the vocalization was emitted. Comparative analyses should also be utilized to determine if differences in vocalizations can be linked to sex, age, and acclimation status.

## 5. Conclusion

For decades, ethological studies have classified animal vocalizations Given that swine are one of the most popular large-animal models used in translational research, it is important to create a catalogue of vocalizations typical to a biomedical laboratory environment. In the present study, perceptual and objective cluster analyses show that the domestic pig vocal repertoire consists primarily of two discrete classes (i.e., low frequency and high frequency) that contain multiple call types. While the vocal classes and call types produced by adult pigs in a biomedical laboratory environment correspond closely to those reported in piglets and wild boars, there are a few call types – including whines, yelps, and stable screams – that are unique to this group of animals. We provided descriptive labels and defining measures for each call type to supply researchers with a common language for future work in pig vocal analysis. As the use of domestic pigs in biomedical research continues to increase, the present results serve as a foundation upon which to build comparative analyses to assess treatment efficacy and improve animal welfare.

## Acknowledgements

The authors would like to acknowledge the efforts of Tapan S. Mehta (biostatistical analysis), Virginia Aida (data collection), & Je’Vana L. Pickens (data collection), Dr. Jeremy Marchant-Forde (concept expertise), Holger Klink (technical expertise) and Ashakur Rahaman (technical expertise).

The authors acknowledge the following financial support for the research, authorship, and/or publication of this article. This research was supported in part by the HEAL Initiative of the National Institutes of Health (NIH) under award number RF1NS135504. This work was also supported in part by a pilot award 121RX00200 from the United States (US) Department of Veterans Affairs (VA), Rehabilitation Research and Development Service. The contents do not represent the views of the NIH or VA or the US Government.

**Acknowledgements**

The authors would like to thank Dr. Jeremy Marchant-Forde for providing expertise in domestic pig behavior and vocal analysis, and Drs. Holger Klink and Ashakur Rahaman for providing guidance in sound analysis methods. The authors would also like to thank the Cornell Lab of Ornithology Sound Analysis and Sound Recording Workshops for their invaluable technical instruction. We thank Dr. Tapan S. Mehta for his assistance and expertise with the statistical analysis. We thank Dr. Virginia Aida and Dr. Je’Vana L. Pickens for their assistance in data collection.

The authors acknowledge the following financial support for the research, authorship, and/or publication of this article. This research was supported in part by the HEAL Initiative of the National Institutes of Health (NIH) under award number RF1NS135504. This work was also supported in part by a pilot award 121RX00200 from the United States (US) Department of Veterans Affairs (VA), Rehabilitation Research and Development Service. The contents do not represent the views of the NIH or VA or the US Government.

